# Driver mutations in GNAQ and GNA11 genes as potential targets for precision immunotherapy in uveal melanoma patients

**DOI:** 10.1101/2022.09.28.509834

**Authors:** Sandra García-Mulero, Roberto Fornelino, Marco Punta, Stefano Lise, Mar Varela, Rafael Moreno, Marcel Costa-Garcia, Alena Gros, Ramón Alemany, Josep María Piulats, Rebeca Sanz-Pamplona

## Abstract

Uveal melanoma (UM) is the most common ocular malignancy in adults. Nearly 95% of UM patients carry the mutually exclusive mutations in the homologous genes GNAQ (aminoacid change Q209L/Q209P) and GNA11 (aminoacid change Q209L). UM is located in an immunosuppressed organ and do not suffer immunoediting. Therefore, we hypothesize that driver mutations in GNAQ/11 genes could be recognized by the immune system. Genomic and transcriptomic data for primary uveal tumors was collected from TCGA-UM dataset (n=80). The immunogenic potential for GNAQ/GNA11 Q209L/Q209P mutations was assessed using a variety of tools and HLA types information. The immune microenvironment was characterized using gene expression data. All prediction tools showed stronger GNAQ/11 Q209L binding to HLA. The immunogenicity analysis revealed that Q209L is likely to be presented by more than 73% of individuals in 1000G database whereas Q209P is only predicted to be presented in 24% of individuals. GNAQ/11 Q209L showed higher likelihood to be presented by HLA-I molecules than almost all driver mutations analyzed. Samples carrying Q209L had a higher immune-reactive phenotype: (i) expression of antigen presenting genes HLA-A (p=0.009) and B2M (p=0.043); (ii) immunophenoscore (p=0.008); (iii) infiltration of immune system cells NK (p=0.002) and CD8+ T lymphocytes (p=0.02). Results suggest a high potential immunogenicity of the GNAQ/11 Q209L variant that could allow the generation of novel therapeutic tools to treat UM like neoantigen vaccinations.

## 1. Introduction

Despite being considered a rare tumor (10 cases per million incidence in Europe), uveal melanoma (UM) is the most common ocular malignancy in adults (1). Prognosis is still poor, with up to 50% of patients developing metastasis, mostly in the liver. Metastatic UM does not have an effective standard treatment available and survival rates have not improved in the last decades (2).

At the molecular level, UM is very different from cutaneous melanoma. Both arise from melanocytes, but they do not share somatic mutations driving carcinogenesis. UM shows exclusive mutations in the GNA gene family. Nearly 95% of UM patients carry the mutually exclusive mutations GNAQ/GNA11 in the hotspot Q209. These mutations change the conserved catalytic glutamine for a Proline, P, or Leucine, L, leading to the constitutive activation of the GTPase domain (3). These oncogenic mutations in G protein-coupled receptor (GPCR) activates pathways including MPAK, PI3K/AKT or YAP/TAZ promoting tumor progression (4).

Unlike cutaneous melanoma, UM do not respond to immune checkpoint inhibitors (5,6). This could be due to several molecular and anatomical differences. UM is located in an immune-privileged organ, protected by the blood-ocular barrier and exhibits an immunosuppressive microenvironment. Because of that, it does not suffer immunoediting (7). Moreover, the tumor mutational burden (TMB) is very high in cutaneous melanoma but low in UM (8). Thus, UM generates low levels of neoantigens and is considered a tumor with low antigenicity (3). Also, we and others showed that immune cell infiltration is associated with poor prognosis in UM (9) (10,11).

Although driver mutations are normally catalogued as non-immunogenic, recent work support the possibility to develop immunotherapeutic drugs against neoantigens derived from recurrent mutations in cancer driver genes in some cases (12). We hypothesize that recurrent mutations in GNAQ and GNA11 genes could elicit T-cell responses. Given the predicted low immune selective pressure in UM, it could represent an attractive target for immunotherapeutic interventions. Also, we hypothesize that different mutations (Q209P or Q209L) could have different antigenicity and response from the immune system. Our objective is to computationally analyze the antigenicity of tumors harbouring GNAQ/11 mutations, to characterize their microenvironment, and to assess their association with clinical phenotypes. Our results suggest that Q209L mutation is more immunogenic than Q209P mutation, irrespectively of the mutated gene (GNAQ or GNA11).

## 2. Methods

### 2.1 Samples

Clinical and mutational data of paired primary uveal tumors and blood samples from patients was collected from TCGA-UM dataset (n=80 pairs). Annotated mutational data was downloaded from the cBioPortal (13). RNA-seq was downloaded in fragments per kilobase per million (FPKM), then converted to log2 scale. Supplementary Table 1 includes a detailed description of patients included in the dataset. Comparison between groups were performed by Chi-squared test for categorical variables and Wilcoxon test for numerical variables. For survival analysis validation dataset, a series of a uveal melanoma from Universitary Hospital of Bellvitge (n=73) with clinical and mutational status information was used.

### 2.2 Immunogenicity prediction of neoantigens GNAQ-L, GNAQ-P and GNA11-L

First, for each mutation, 19 mers amino acid sequences were extracted using an in-house script. Mutated amino acids were in the center of the sequence. Wild type sequences were also generated. The immunogenic potential for GNAQ-L, GNAQ-P and GNA11-L was assessed in a variety of binding prediction tools (NetMHC, NetMHCpan, NetMHCcons, NetMHCpanstab, MHCSeqNet and MHCflurry) using HLA supertypes and all 9mer combinations from the two mutated sequences as input (14–21).

Apart from solo binding prediction, NetCTL tool was used to predict proteasomal C terminal cleavage and TAP transport efficiency (22). The proteasome cleavage event is predicted using the version of the NetChop neural networks trained on C terminals of known CTL epitopes as described for the NetChop-3.0 server (23). The TAP transport efficiency is predicted using the weight matrix-based method described by Peters et al (22). NetCTL predicts MHC peptide binding using neural networks in NetMHC server and then calculates a combined score for the three measures. As an input, fasta files with GNAQ and GNA11 protein sequence was used.

### 2.3 HLA presentation scores

All HLA-presentation scores were defined starting from eluted ligand likelihood percentile ranks of peptides with respect to HLA allotypes obtained from the NetMHCpan-4.0 prediction method (15). NetMHCpanI were run (HLA type I only predictions) on all neopeptides of length 8 to 11 generated by each of the 3 mutations (GNAQ-L, GNAQ-P and GNA11-L) against a set of 195 HLA(-A/-B/-C) types found in the >1,000 individuals of the 1000Genomes project. For each individual there was information about 6 HLA types.

Each mutation was mapped to a protein sequence and associated to a set of 38 mutated peptides using an in-house Python script to generate all possible peptides of length 8 to 11 that spanned the mutation. A wild type peptide was associated to each specific mutant peptide that was identical to the mutant peptide except that the mutated amino acid is reverted to the wild type one. For each peptide in this set, the program NetMHCpan-4.0 (57) was used to calculate the eluted ligand likelihood percentile rank and predict the interaction core peptide (Icore) with respect to all HLA allotypes. The elution rank takes values in the range from 0 to 100, with lower values representing higher presentation likelihoods. We defined the presentation score of a mutation with respect to a specific HLA allotype as the minimum elution rank among all associated peptides but excluding those with a wild type Icore. We called this presentation BR score.

PHBR score (Patient Harmonic-Mean Best Rank) was calculated by combining the six best rank socres of the six HLA allotypes using a harmonic mean. Also, we calculated our Population-Wide Median Harmonic-Mean Best Rank (PMHBR) as the median of the PHBR scores of a mutation calculated over a set of individuals. Lower PMHBR scores correspond to higher likelihoods for the mutation to be presented across our 1000G or TCGA populations (24).

### 2.4 Immune microenvironment characterization

The immune microenvironment of the samples was characterised using gene expression data and a variety of bioinformatics tools. The immunophenoscore (IPS) function was used to measure the immune state of the samples by the quantification of four different immune phenotypes in a given tumor sample (Antigen Presentation, Effector Cells, Suppressor Cells and Checkpoint markers), using gene markers. Also, it computes an aggregated z-score summarizing the four immune phenotypes (26). Finally, samples were scored using the gene set variation analysis (GSVA) method with 18 gene markers lists from ConsensusTME (27) and the T-cell inflammatory (TIS) signature (28).

### 2.5 Survival analysis

A survival analysis was done with a cohort of patients from the Bellvitge University Hospital (n=73). Progression free survival (PFS) was assessed between patients harboring Q209P (n=25) and Q209L (n=48) mutation. Kaplan–Meier curves were plotted to represent the result and log-rank test was computed.

## 3. Results

### 3.1 GNAQ/GNA11 mutations in TCGA-UM dataset

GNAQ and GNA11 were the most frequent missense mutations in TCGA-UM dataset and were mutually exclusive (**Figure 1A**). Out of 80 TCGA-UM patients, 34 patients carried GNA11 p.Q209L (hereafter GNA11-L), 10 patients carried GNAQ p.Q209L (hereafter GNAQ-L), and 27 patients GNAQ p.Q209P (hereafter GNAQ-P). The other 9 samples were wild type at the position of interest; two patients carried GNAQ p.R183Q mutation, one more patient carried GNAQ p.G48V, one patient GNA11 p.R183C, and one patient GNA11 p.R166H. Two individuals were mutant at the same time for GNAQ and GNA11 but the second hit was not in position 209 (one case at positions GNAQ p.Q209L and GNA11 p.R166H; second case at positions GNAQ p.R183Q and GNA11 p.R183C) (**Figure 1B**).

**Figure 1.**
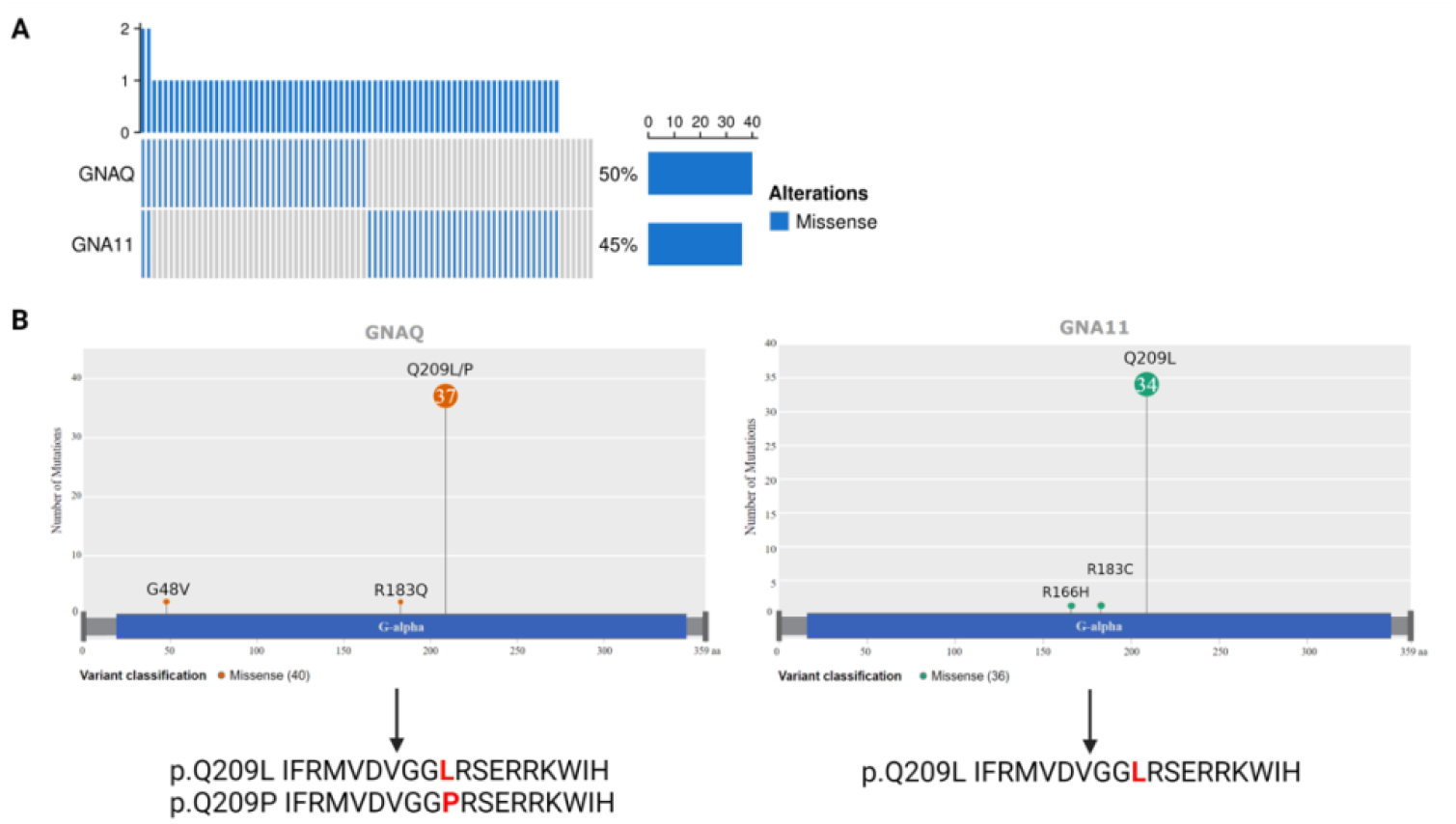
GNAQ and GNA11 mutations in TCGA UM samples. **A**) Mutational status of GNAQ and GNA11 genes. Barplot shows mutated patients in blue and wild type in grey. Frequency of alterations are 50% for GNAQ and 45% for GNA11. **B**) Lollipop plot showing GNAQ and GNA11 mutations across the proteins and resulting peptides harbouring Q209P and Q209L mutations. Aminoacidic changes are marked in red.

Despite being located in different chromosomes (Chr. 9 and Chr. 19, respectively), GNAQ and GNA11 genes are highly homologous and so are the resulting proteins. A BLAST alignment showed 90% identity between the two proteins (**Supplementary Figure 1**). GNAQ-L and GNA11-L suffered the same amino acid change in position 209 (from Q -Glutamine-to L -Leucine-), and given the high homology between these two proteins, the resulting 19-mer peptide in which the mutation is centred were identical. On the other hand, GNAQ-P changed from Q (Glutamine) to P (Proline). Because of this, and since we planned to study the potential immunogenicity of those mutations rather than protein function, we decided to compare patients harboring P mutated vs. patients harboring L mutated, irrespectively of the gene of origin (**Figure 1B**). In total, 71 (89%) patients carried the Q209P/L amino acid change, of which 44 (62%) carry amino acid change p.Q209L and 27 (38%) carry change p.Q209P.

To see whether there was any association between the different change Q209P or Q209L and the different clinical variables in the dataset, we performed a statistical test by mutation change (**Supplementary Table 2**). No association was found with age, sex, overall survival time and status, progression free survival status, recurrence, fraction of genome altered, SCNA subtype cluster, BAP1 mutation, Chromosome 3 status (disomy or monosomy), Chromosome 8 status (disomy or polysomy) or immune cluster. The only significant association was the mutation count (Wilcoxon test, p-val=0.028), indicating that patients with Q209L mutations have a slightly higher number of mutations (a mean of 13.3 vs. 11.1). However, this is not significant when multitesting correction was applied.

### 3.2 Binding affinity prediction of neoantigens GNAQ-L, GNAQ-P and GNA11-L

The 19 length peptides for GNAQ-L/GNA11-L (Q209L) (IFRMVDVGGLRSERRKWIH) and GNAQ-P (Q209L) (IFRMVDVGGPRSERRKWIH) were used to test the antigenicity of these mutations using a total of 7 different binding prediction tools, to avoid any bias related to similar Machine Learning algorithms or datasets used for the training. Most of these prediction tools focus on scoring the affinity of the inputted peptides for a specific HLA. However, NetMHCstabpan, which calculates a combined score for the affinity and stability of the binding was also used. As input, tools used all possible 9mer combinations from the two 19mer mutated sequences studied. Apart from the peptide sequences, we used all the HLA superfamilies for the predictions.

The outputs of the different tools were diverse. The NetMHC tools and MHCflurry calculate an affinity value measured in nM, which is used to filter the binders or no binders. These affinity values are also shown as a logaritmic tranformations, called %Rank. Only the 9mers with a value of 500nM and below are considered binders. On the other hand, the output of MHCSeqNet is a probability value between 0.0 and 1.0, where 0.0 refers to a non-binder and 1.0 to a strong binder. Only those with more than 60% probability of binding were taken. Lastly, MixMHCpred does not provide affinity value, instead, it calculates a Score and a %Rank value for each HLA allele. For a single allele, scores larger than 0 correspond to %rank smaller than 1%. Therefore, in the case of this tool, we only choose the 9mers in which the best allele score is higher than 0 (**Supplementary Table 3**).

In summary, all methods predicted Q209L mutation as being more inmunogenic (assuming the higher binding values the more immunogenic) than Q209P, except for NetMHCPan that predicted equal number of peptides with binding affinity (**Figure 2**). A total of 12 non-unique bindings with different HLA types were found for Q209P variant whereas a total of 29 bindings were found for Q209L. Although only 4 out of the 7 tested tools give information about the strength of the binding, no Q209P neoantigen was predicted as strong binder. However, 5 putative neoantigens were classified as strong binders in the case of Q209L (**Table 1)**. The HLA haplotypes giving rise to strong bindings with Q209L mutation were HLA-A*03:01, HLA-B*27:05 and HLA-B*39:01. For Q209P mutation, the HLA with strong bindings were HLA-A*03:01 and HLA-B*07:02 (**Supplementary Table 3**).

**Figure 2.**
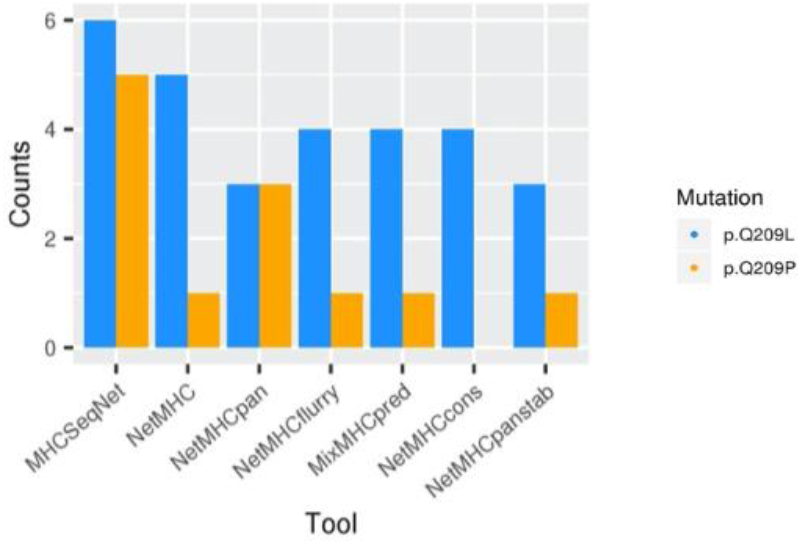
HLA-Q209L and HLA-Q209P binding prediction. cross seven prediction tools (in x axes), using HLA supertypes genotypes.

**Table 1.** Number of predicted bindings. WB: weak binding, SB: strong binding.

| Mutation | NetMHC | NetMHC-pan | NetMHC-cons | NetMHC-stabpan | MHC-SeqNet | MHC-flurry | MixMHCp red | Total |
| --- | --- | --- | --- | --- | --- | --- | --- | --- |
| Q209P | 1 (WB) | 3 (WB) | 0 | 1 (WB) | 5 | 1 | 1 | 12 |
| Q209L | 5 (3 WB + 2 SB) | 3 (2 WB + 1SB) | 4 (2 WB + 2 SB) | 3 (WB) | 6 | 4 | 4 | 29 |

To know if these mutations would be likely to be presented by any individual from the 1000Genomes database, as a sample of healthy population, we calculated how many individuals have at least one mutant peptide (length 8 to 11) that has presentation likelihood below a given threshold for at least one of the HLA types of the individual. For threshold % rank <0.5 (Strong binding), up to 73% of individuals were predicted to present Q209L peptide, while only 24% of individuals were predicted to present Q209P peptide.

Looking at threshold %rank <2 (Weak binding), 88% of individuals were presenting Q209L peptides and 74% of individuals present Q209P. Moreover, we generated a BR score for each sample carrying Q209L by taking the minimum BR score of all 6 BR per patient. The 69.7% of samples have at least one strong binding (BR<0.5), while 16.3% have a weak binding (0.5<BR<2) and 14% have no binding (BR>2).

Next, the percentage rank score of mutant peptides were compared to the percentage rank of their corresponding wild type (WT) peptides. This may be relevant because given the similarity between WT and mutants (a single aa difference) it is possible that if the WT is presented, the mutant (even if presented) may be subjected to tolerance mechanisms and thus not be immunogenic. For % rank < 0.5 threshold, in 59% of individuals the mutated peptide Q209L is predicted to presented with strong binding while the Q209L WT is not. On the other side, only 8% of individuals are predicted to present the Q209P mutated peptides and not the Q209P WT peptide. So, mutation Q209L has the most encouraging differences with respect to WT.

Finally, the HLA binding affinity was predicted through a score of antigenicity for the two mutations Q209P and Q209L. This score is calculated based on the “Best rank” score of NetMHCpanI for the 100Genomes population. As explained, the BR score is the Best Rank for each individual, while the PMHBR is the median population BR score. The PMHBR score of Q209P is 3.66, while the PMHBR score of Q209L is 0.62. Then, we have compared these scores to other driver mutations, and we see that Q209L mutation has one of the lowest scores, meaning that it has higher likelihood to be presented across the population than most of the driver mutations of different cancer types (**Figure 3**).

**Figure 3.**
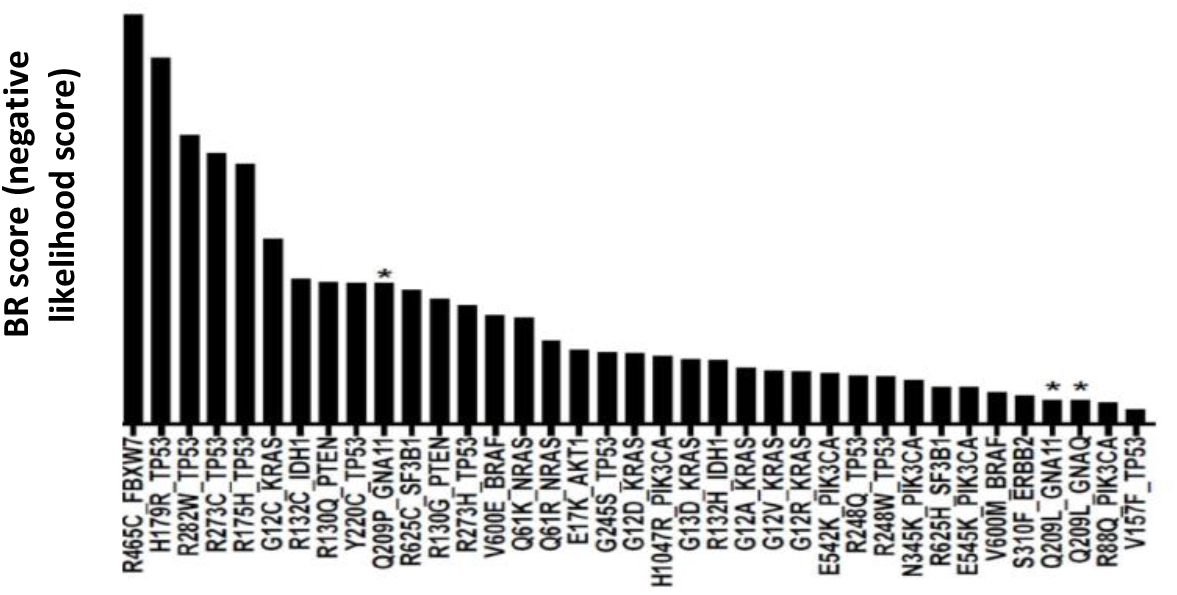
PMHBR score of a list of driver mutations across 1000 Genomes individuals. The lower PBHBR score the higher probability to be presented. Q209L shows higher likelihood to be presented by HLA molecules than Q209P and most driver mutations in cancer. Asterisks marks Q209P and Q209 L mutations.

Apart from binding to HLA, for a neoantigen to be presented it needs to be processed by the proteasome and transported by the TAP mechanism. We used NetCTL to predict proteasomal C terminal cleavage and TAP transport efficiency. As a result, for Q209L we got 3 putative neoantigens whereas we got only 2 in the case of the Q209P. For Q209L, NetCTL selected 9 mer FRMVDVGGL as a good candidate to be presented by HLA-B*27:05 and HLA-B*39:01 and RMVDVGGLR to be presented by HLA-A*03:01. These two peptides were also predicted to be binders by all the other tools. The former as a strong binder and the later a weak binder.

Taking together, all these results points to Q209L mutation to be more immunogenic, being predicted to be properly processed and presented with good affinity and stability.

### 3.3 HLA haplotypes frequencies with uveal melanoma risk and survival

Next, we wanted to assess if having different HLA haplotypes (implying different binding affinity for Q209L) has an impact on uveal melanoma risk or survival. First, we wonder if there was a relationship between HLA haplotype frequency and the BR scores. In the general population, the BR score of Q209L mutation is not correlated with HLA frequency for HLA-A and HLA-B genes (**Figure 4A**), while BR score and HLA-C exhibited a non-significant trend towards negative correlation. For UM patients, the negative correlation between HLA-C haplotype frequency and the BR score is stronger (Spearman correlation, p=0.057, **Figure 4B**). Results from 1000G population pointed to HLA-C*07:02 as the allele with the more frequency and lower BR score. On the contrary, HLA-A*24:02 is an example of frequent allele with no predicted binding affinity for Q209L (**Figure 4C**).

**Figure 4.**
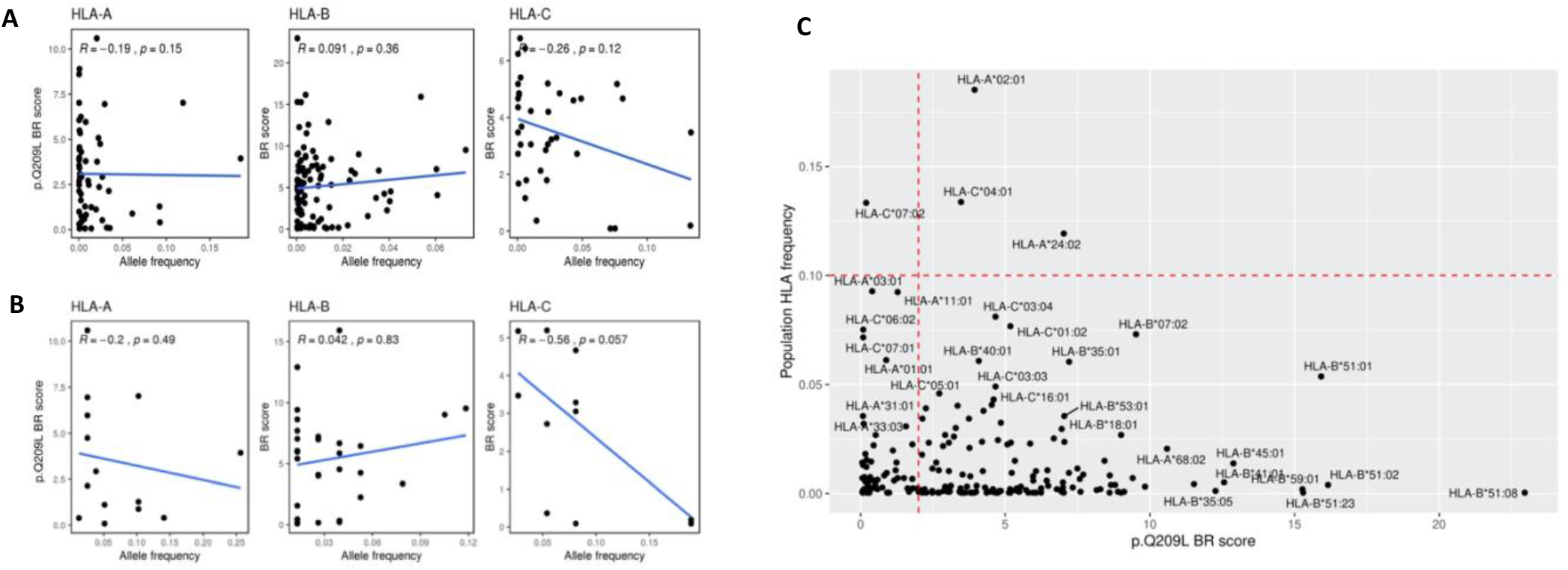
Correlation between presentation probability and HLA haplotypes. Plots showed correlation between BR score of Q209L and the HLA-A, HLA-B and HLA-C type frequency in **(A)** general population and **(B)** in UM patients. **C)** Plot showing BR score for Q209L in x axis and population HLA frequency in y axis with HLA haplotypes of high frequency annotated.

HLA frequencies between uveal patients and general, healthy population (1000G) were compared by binomial test and resulting frequencies were plotted in a radar plot (**Figure 5A, Supplementary Table 4**). As a result, 10 haplotypes showed differences at FDR<0.05 between uveal and population frequency, of which 9 showed higher frequency in uveal melanoma patients; HLA-A*01:01, HLA-A*02:01, HLA-B*08:01, HLA-B*15:01, HLA-B*18:01, HLA-B*44:02, HLA-C*01:02, HLA-C*05:01, HLA-C*07:01, HLA-C*12:03. Of those alleles, only HLA-C*07:01 and HLA-A01:01 have BR score of high antigenicity (BR<2), while the other seven with higher frequency in uveal melanoma have high BR value scores (BR>2; low antigenicity scores), suggesting a genetic selection in uveal melanoma patients made neoantigen Q209L to be hidden.

**Figure 5.**
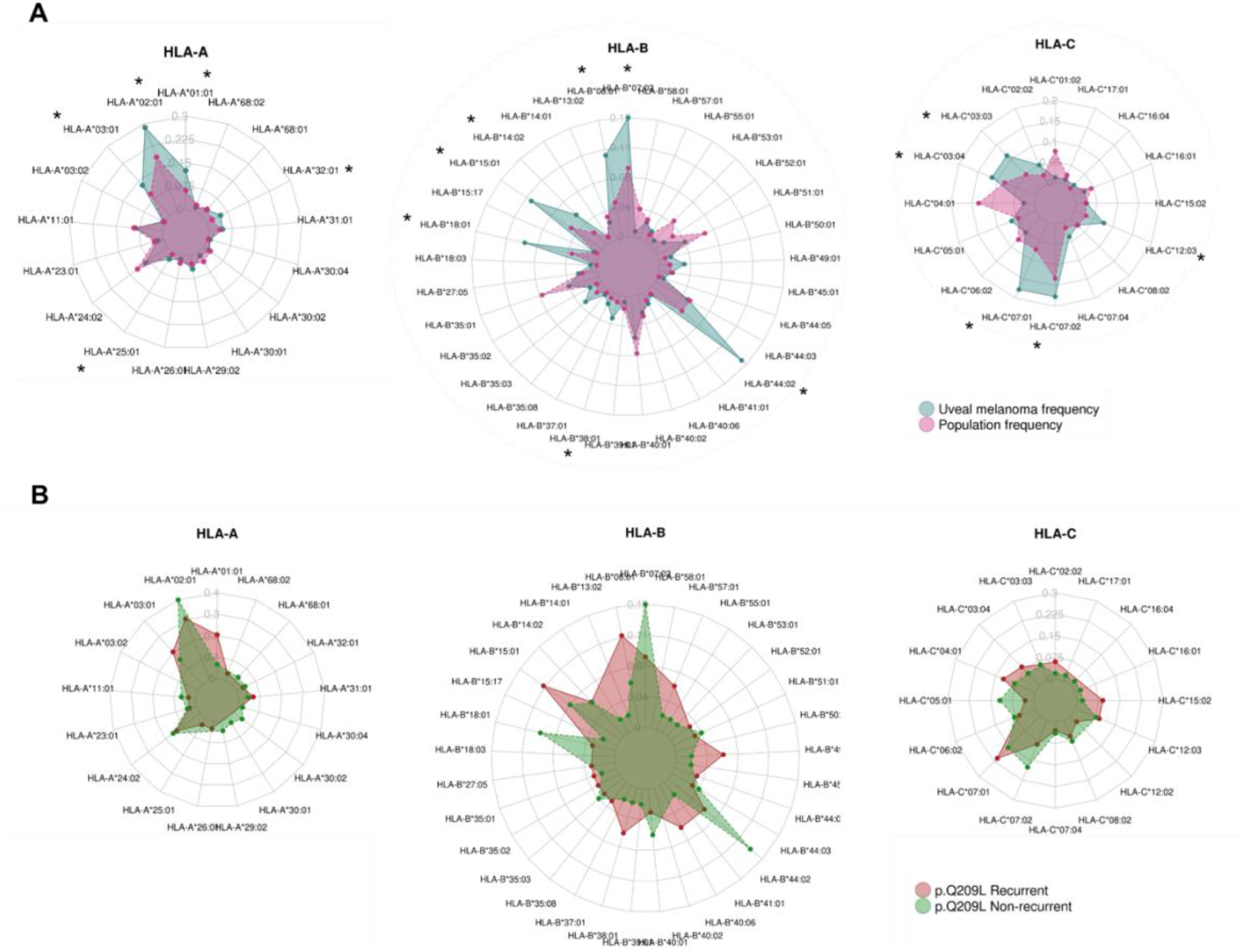
HLA frequency association with UM risk and survival. Radar plots comparing frequencies in HLA haplotype for HLA-A gene, HLA-B gene and HLA-C genes **A**) Between UM patients (green) and 1000 Genomes individuals (purple), and between recurrent (red) and non-recurrent (green) patients harbouring Q209L (**B**). Asterisks correspond to haplotypes with statistical differences by Binomial test (FDR p-adjusted < 0.05). Only haplotypes which are present in uveal melanoma patients are depicted.

The same analysis was performed for comparing the HLA frequencies between patients harbouring Q209L or Q209P mutations (**Supplementary Figure 2**). In this case, no statistical differences were found between the frequencies. Finally, to find out whether there could be selection towards lower antigenic binding in patients carrying the highly antigenic Q209L change and relapsing, we compared the HLA frequencies in patients carrying Q209L mutation, between recurrent and non-recurrent uveal melanoma samples. As in the previous comparison, none of the HLA haplotypes compared by binomial test showed statistically significant differences. On the contrary, there is a tendency towards higher frequency in HLA-B*44:02, HLA-B*07:02 and HLA-B*18:01 in non-recurrent samples, which are three haplotypes with low binding affinity to Q209L. (**Figure 5**). Also, we wonder if HLA haplotypes with higher chances of presenting Q209L were absent in uveal melanoma patients. However, there are not statistically significant differences in BR score between haplotypes present and missing in uveal melanoma patients (**Supplementary Figure 3**).

Finally, a survival analysis was done in a total of 73 human samples from the Bellvitge University Hospital (n=73) between Q209L and Q209P. As a result, Q209L patients showed slightly better progression free survival (PFS) than Q209P patients (Log-rank test p=0.038). (**Supplementary Figure 4**). However, in TCGA data, we have not found any relationship between P/L mutations and prognosis.

In summary, no clear associations have been found between HLA haplotypes and risk of suffering uveal melanoma neither between HLA frequency and survival. It is important to point out that there is a possibility that we do not find statistical differences between recurrent (n=17) and non-recurrent (n=26) Q209L patients due to the low sample size.

### 3.4 Samples harboring GNAQ-P or GNA-L mutations showed differences in the tumor microenvironment

We used expression data to characterize the immune state of samples carrying Q209L mutation or Q209P mutation. First, we evaluated whether there were differences in the levels of antigen processing and presentation genes (**Figure 6A**). All genes related to MHC class I showed higher gene expression in patients carrying Q209L mutation (Wilcoxon test; HLA-A, p=0.009; HLA-B, p=0.039; HLA-C, p=0.034, B2M, p=0.043).

**Figure 6.**
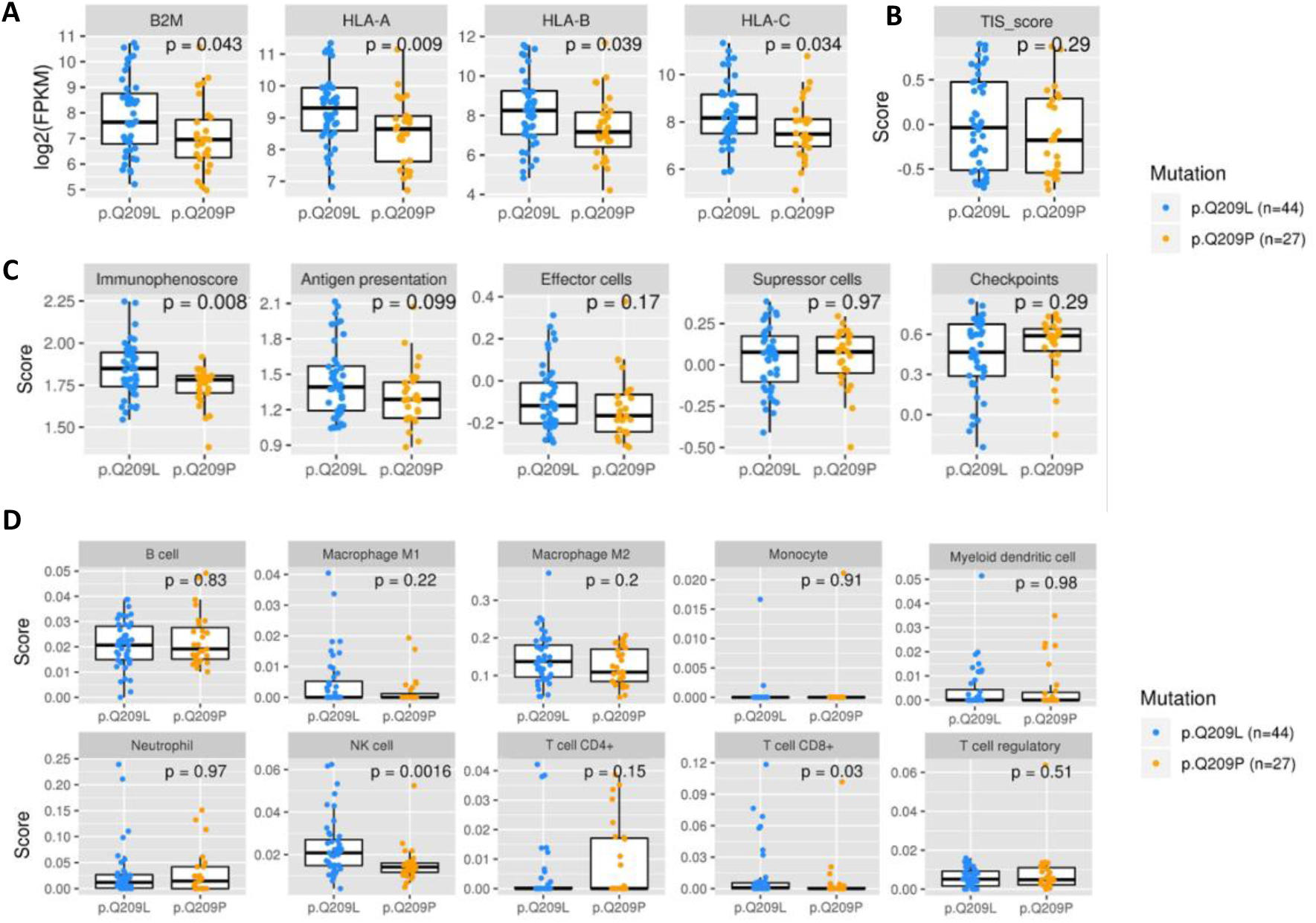
Characterization of immune state in patients carrying Q209L variant and Q209P variant. **A**) Levels of expression of antigen presenting genes B2M, HLA-A, HLA-B and HLA-C. **B**) T cell Inflammatory signalling (TIS) score. **C**) Immunophenoscore (IPS), antigen presentation, effector cells, suppressor cells, and checkpoints scores. **D**) Immune cell infiltration. Wilcoxon test were used to calculate statistically significant differences (p-value < 0.05).

Next, we used a number of tools to characterize the immune system activation status of samples. The T-cell inflamed signature (TIS score) was estimated and showed no differences between Q209L and Q209P mutated patients (**Figure 6B**). The Immunophenoscore, that is used as global score of the immune state of the samples, was significantly higher in Q209L mutated patients (p=0.0081) (**Figure 6C**). This score is based on four sub-scores that represent the activation of Antigen presentation, Effector cells, suppressor cells and checkpoint markers (neither of those showed statistically significant differences, although there is a tendency to higher antigen presentation and effector cells activation in Q209L patients).

To explore the infiltrate in detail, we used Quantiseq method for estimating the infiltration of immune cells in the tumour microenvironment (**Figure 6D**). We found higher infiltration of T cells CD8+ (p=0.03) and NK cells (p=0.0016) in Q209L patients. To validate these results, we estimated the scores with a second method, called ConsensusTME (consensus tumor microenvironment) (**Supplementary Figure 5**). In agreement with the previous method, we found that patients carrying Q209L mutations tended to higher infiltration scores for CD8 T cells (p=0.065). In contrast, we found no differences in NK cells. No differences were found for the other cell types with this method, although there was a trend towards higher scores of B cells in Q209P patients. Despite the variability between the methods, all results suggest a distinct immune microenvironment modulation, indicating a high immune reactive phenotype in tumours harbouring Q209L mutations.

Finally, to look at differences in activated biological pathways, a differential expression followed by gene-set enrichment analysis between Q209L carriers and Q209P carriers was performed. A total of 12 genes were found at p-value<0.05 and absolute values of logFC>1, of which 9 were overexpressed in Q209L patients and 3 were overexpressed in Q209P patients (**Supplementary Table 5**). In the functional analysis, as expected, most enriched gene sets for Q209L patients were related with immune system (IFN-γ, p-adj=1.38e-12; IFN-α, p-adj=3.06e-8, IL6/JAK/STAT3, p-adj=1.4e-3). Also, other pathways related with tumour growth and metabolism emerged (mTOR signalling, p-adj=5.16e-5; hypoxia, p-adj=0.011, oxidative phosphorylation, p-adj=0.03, and fatty acid metabolism, p=0.04) (**Supplementary Figure 6, Supplementary Table 6**). Otherwise, there was not any pathways enriched in Q209P patients. This result suggests a crosstalk between immune infiltrate and other components of the tumour biology in Q209L carriers.

## 4. Discussion

Activating mutations in the Gαq signaling pathway at the level of GNAQ and GNA11 genes are considered alterations driving proliferation in UM. Lot of research has been devoted to understanding molecular mechanisms behind these alterations, which transfer signaling from GPCRs to downstream effectors by activating the pathway constitutively. Also, to develop blocking drugs (29). Despite these efforts, no novel treatment targeting this pathway has improved the prognosis of UM patients. Due to the exclusive immune microenvironment of UM, here we propose to study these driver mutations from an immunogenic point of view.

Our hypothesis is that different amino acids in the same position (P or L) activates different immune response in the patient, rather than being GNAQ or GNA11 mutant. To the best of our knowledge, little is known about differences between tumors harboring Q209P or Q209L mutations. A study by Maziarz et al showed fundamental difference in the molecular properties of Gq Q209P compared with proteins harboring Q209L, due to different structural conformations of the aberrant proteins (30).

Contrary to other driver mutations such as those in p53 or BRAF, among others; GNAQ and GNA11 are UM-exclusive mutations. On one hand, these alterations could help cancer cells to acquire an eye-specific adaptation. On the other hand, it might be hypothesized that tumoral cells harboring these mutations in other organs are destroyed by immune system in early stages of the disease. In this regard, it has been reported that highly recurrent oncogenic mutations have poor HLA class I presentation (31). Punta et al reported that the median PMHBR of highly recurrent driver mutations in TCGA is 1.84 whereas the median PMHBR of passenger mutations in TCGA is 1.391. Thus, a driver mutation’s frequency in cancer patients negatively correlates with the population ability to present it (24,31). Our results point to Q209L to be more immunogenic that Q209P in 1000G population. Despite being a driver mutation, it was more likely to be presented in comparison with other recurrent ones. In agreement, all tested tools except NetMHCPan predicted Q209L derived peptide as high immunogenic.

Neoantigens shared among groups of patients have become increasingly popular therapeutic targets. Obviously, non-recurrent, passenger mutations generating neoantigens needs personalized logistics to be therapeutically exploited. On the contrary, public mutations simplifies all this process. In this regard, several public neoantigens from mutations in KRAS, BRAF and TP53 genes has been described so far (12). It is worth to mention a recent work by Samuels et al. describing the combination of HLA-A*01:01 and driver mutation RAS.Q61K as potentially immunogenic in 3% of melanoma patients (39).

We have found differences in immune system activation and infiltration between Q209L and Q209P tumors, being Q209L those scoring better in immunophenoscore. In agreement, Q209L tumors showed higher expression of genes related to antigen presentation. Interestingly, Q209L tumors showed higher infiltration of T-cells and NK cells. It has been reported that normal ocular cells express little or no MHC class I molecules in order to avoid recognition by cytotoxic T-cells. Aqueous humor or eye contains immunosuppressive factors inhibiting NK cells such as TGF-beta or MIF. Paradoxically, metastasizing cells in UM upregulate HLA molecules. Probably, this is because uveal melanoma cells with lower HLA expression are susceptible to be detected and eliminated by NK cells (32). In agreement, in-vitro studies have demonstrated the ability of cytotoxic NK cells to detect and kill uveal melanoma cells (33,34).

Also, differences at functional level have been found. Interestingly, Q209L score better in pathways related to inflammation like interferon alpha and gamma response reinforcing those tumors to be more immunogenic. However, no changes in the inflammatory microenvironment neither in HLA expression has been found in a similar study comparing Q209L vs. Q209P primary uveal tumors (35).

Although a trend was observed towards more frequent HLA in UM showing low BR scores, no clear associations have been found between HLA haplotypes and risk of suffering uveal melanoma neither between HLA frequency and survival. These suggest that genetics of patients do not impact directly on disease initiation or progression through Q209L presentation, or at least there are other implicated factors. One could expect HLA alleles showing low BR score (meaning high likelihood of the neoantigen to be presented by HLA) in healthy population and HLA alleles showing high BR score in UM patients. The low number of UM samples prevented us to discard the hypothesis that people presenting Q209L neoantigen are at lower risk to develop UM. Interestingly, a negative correlation has been found between BR score and HLA-C frequency in both uveal patients and general population suggesting HLA-C as the best presenting allele for this specific neoantigen.

In terms of prognosis, mutations in GNA11 have been moderately associated with poor prognosis and found more frequently in metastatic UM; in comparison with GNAQ mutations (36,37). Other analysis, however, found not differences (35). Looking at amino acidic change, in TCGA-UM data, a marginal p-value of 0,06 pointed to Q209L to be associated with high risk of relapse. No differences in survival status were found. However, in controversy, our results in an independent dataset of primary UM samples showed Q209P patients to have poor prognosis (log-rank=0,04). Interestingly, Terai et al. identified that differences in mutation patterns (Q209P vs. Q209L) in GNAQ and GNA11, rather than GNAQ and GNA11 themselves, might predict the survival of metastatic UM patients. After development of metastasis, patients with GNAQ Q209P mutant tumors had a more favorable outcome than patients with GNA11 Q209L and GNAQ Q 209L mutant tumors (38). Also controversial, but in the primary tumor setting, a work by van Weeghel et al found not differences in prognosis based on Q209P or L mutation but in Chromosome 3 status (monosomy or disomy), as previously reported. In our data, there is not association between Chromosome 3 status and Q209P or L mutation.

This study has several limitations. It has not been validated in independent datasets because of scarce data about amino acidic change in GNAQ, GNA11 mutations. Functional analysis comparing tumors harboring Q209P and Q209L could be biased by differences in number of samples between the two groups. Unfortunately, binding predictors do not perform well with HLA-II so the role of these genes deserves further study. Also, prediction binding algorithms could produce false positive results. The limited sample size is also a drawback. Finally, the study is primarily computational.

Despite the shortcomings, it is worth to mention that an existing patent (WO2019241666) validates our observations. It already defines a technology for the development of a vaccine to treat uveal melanoma based on GANQ/GNAL11 mutations. It shows how the binding of the mutated peptide FRMVDVGGL, which was also found in our study, with the HLA is more immunogenic than the binding with wild type peptide. Also, they describe that the critical amino acids for the binding were R in position 2 and Q/L in position 9, located in MHC pocket acting as an anchor.

Treatment of UM continues to be a challenge, especially in metastatic patients. Although preliminary, our work paves the way for future therapeutic options such as NK cell therapy or neoantigen vaccines. In this study, we report that GANQ/GNAL11 mutations can generate immunogenigity and we have proposed a potential candidate for neoantigen vaccine targeting uveal melanoma.

## Acknowledgements

We thank CERCA Program, Generalitat de Catalunya for institutional support. We also thank the Consortium for Biomedical Research in Epidemiology and Public Health (CIBERESP).

## Supplementary material

**Supplementary Figure 1.**
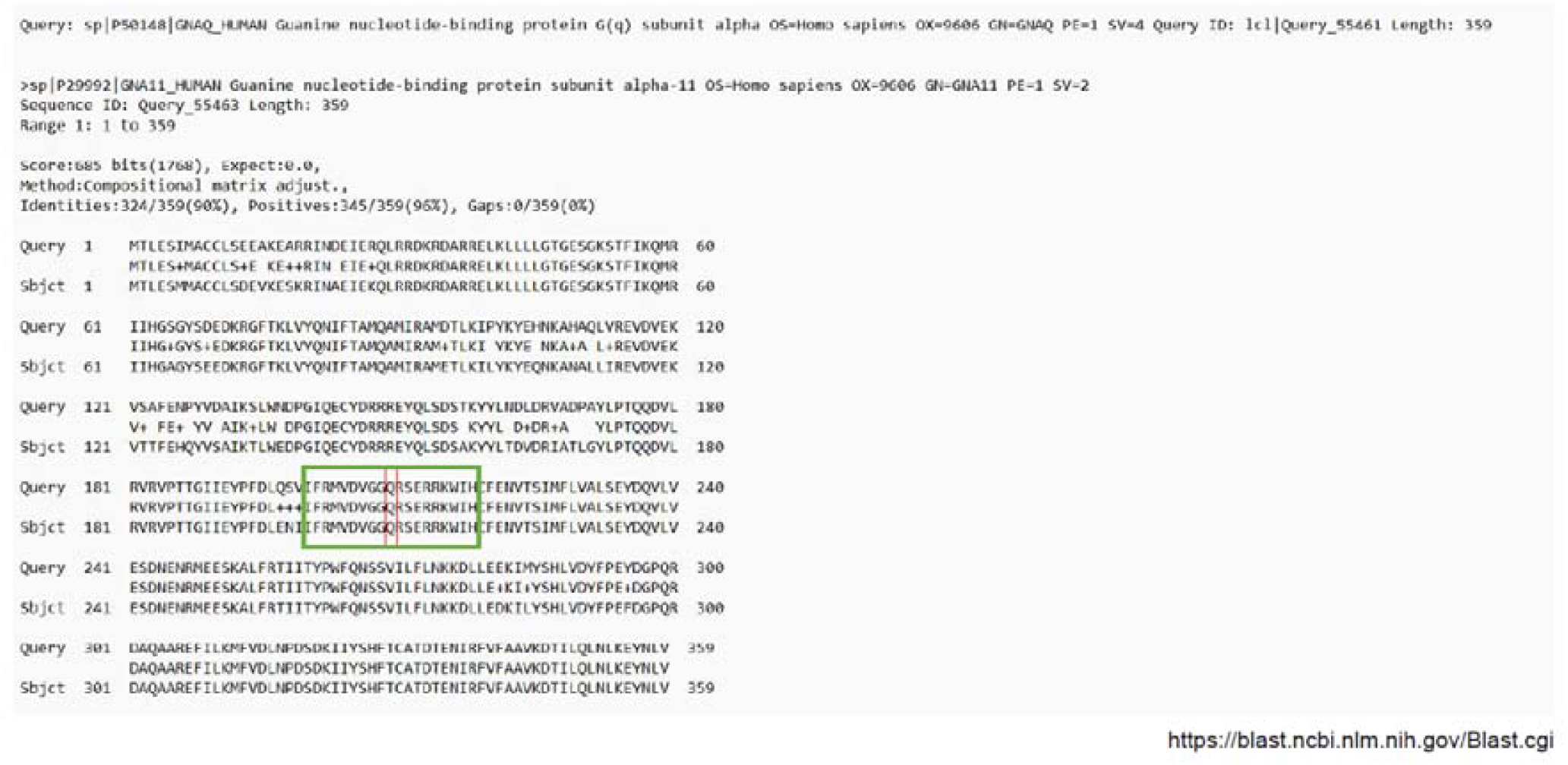
GNAQ and GNA11 protein alignment. Green square marks selected peptides for subsequent binding prediction analysis. Q Amino acid changing to L or P in mutated proteins are marked in red.

**Supplementary Figure 2.**
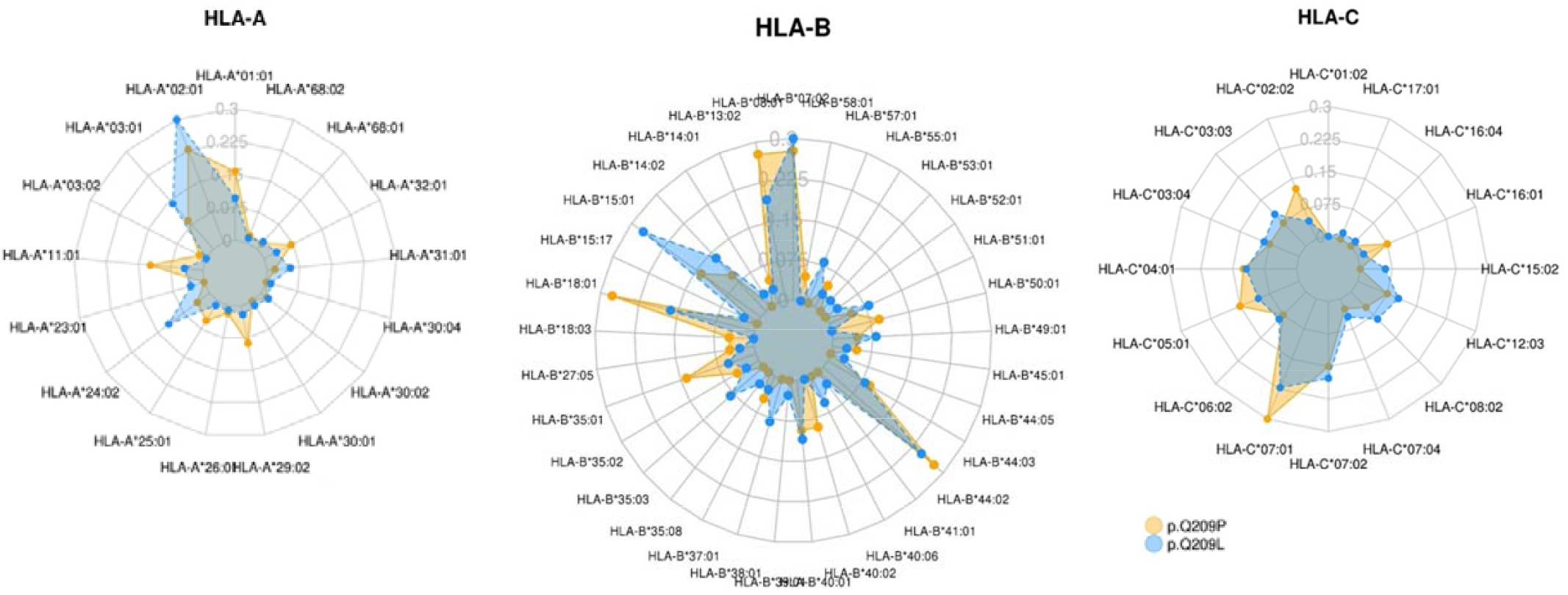
Radar plots comparing frequencies in HLA haplotype for HLA-A gene, HLA-B gene and HLA-C genes between patients harbouring Q209L and Q209P. Asterisks correspond to haplotypes with statistical differences by Binomial test (FDR p-adjusted < 0.05).

**Supplementary Figure 3.**
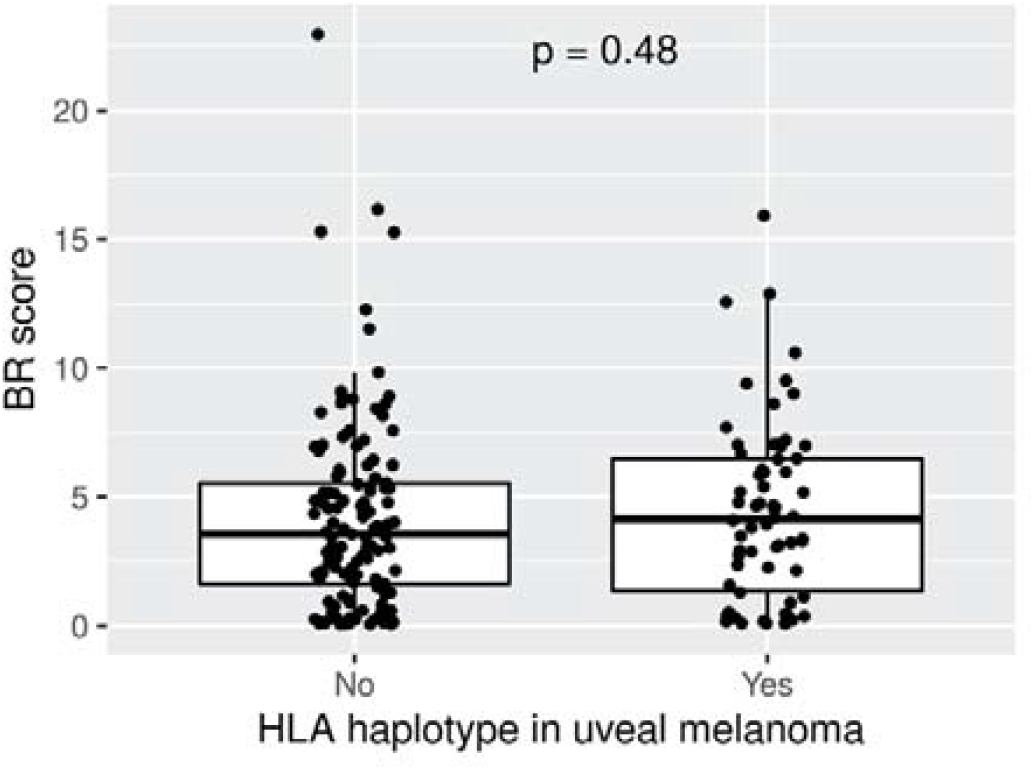
Comparison of BR scores between present and absent haplotypes in uveal melanoma patients. Not statistically significant differences were observed.

**Supplementary Figure 4.**
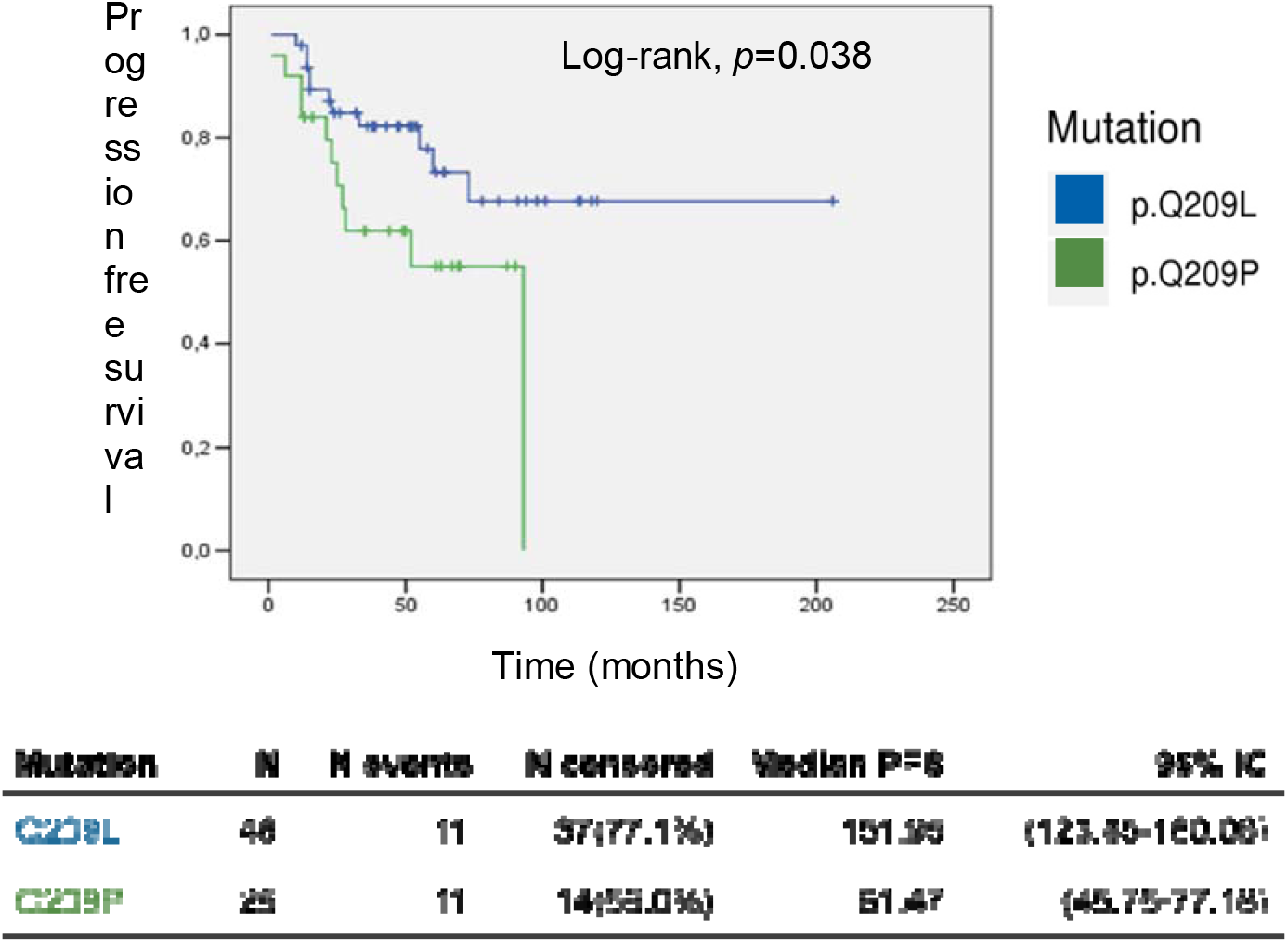
Survival analysis. Kaplan-Meier curve showing Q209L carriers having better disease-free survival (DFS) than Q209P carriers.

**Supplementary Figure 5.**
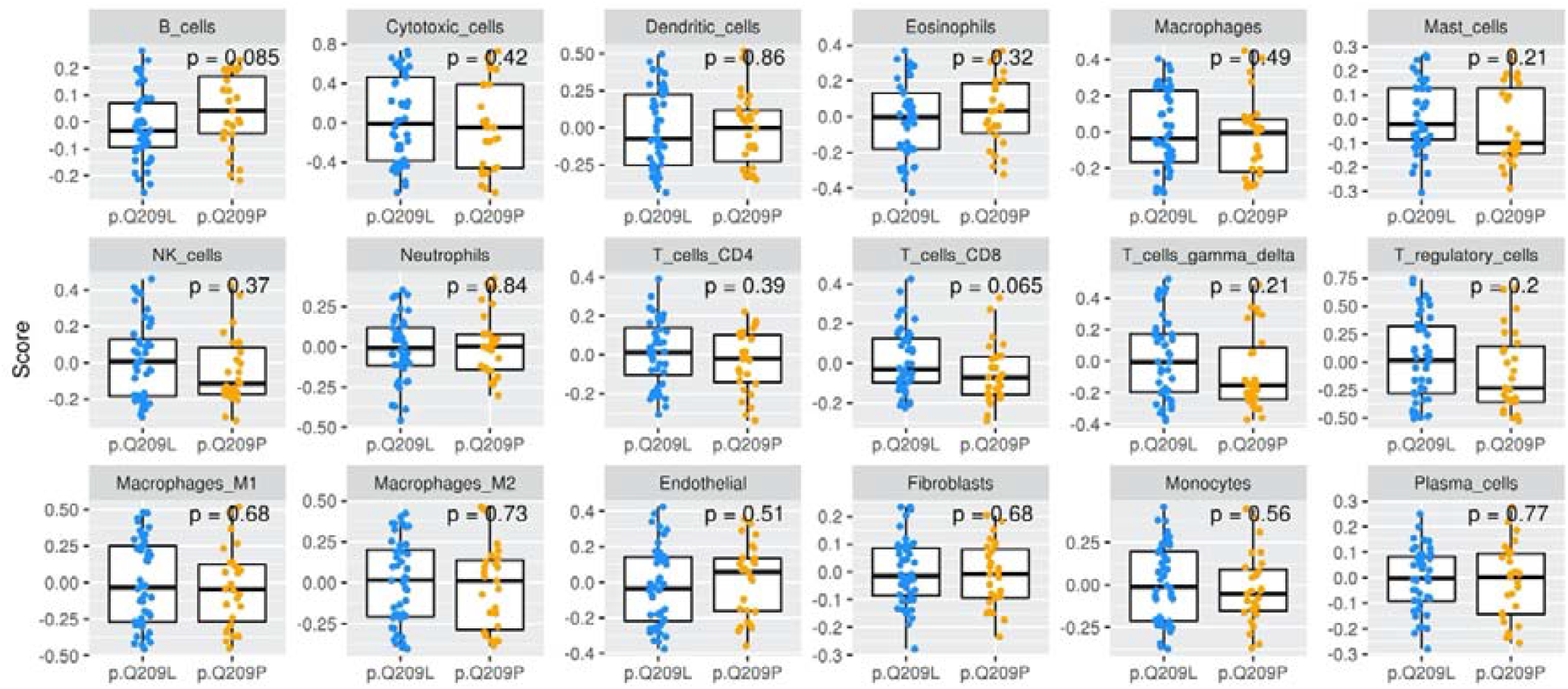
Cell infiltration using ConsensusTME tool.

**Supplementary Figure 6.**
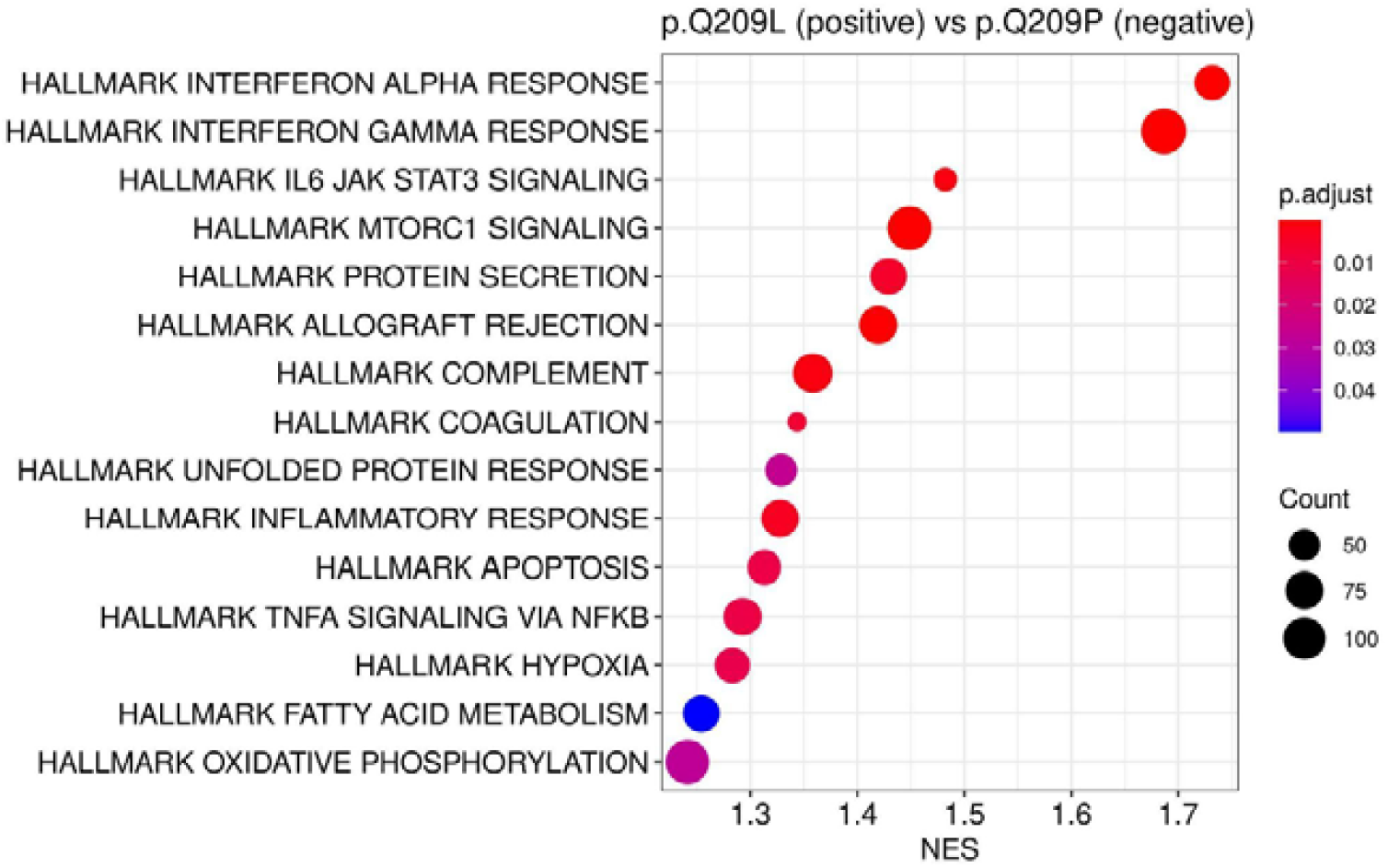
Functional enrichment analysis. Plot showing statistically significant functions over-expressed in in samples carrying Q209L variant. Gene Set Enrichment Analysis (GSEA) sere used. amino-acid change. P-values for categorical variables were calculated by Chi-Squared Tests. P-values for continuous variables were calculated by Wilcoxon tests.

**Supplementary Table 1.** Baseline characteristics of 80 TCGA-UM samples

|  | *[ALL]* | *N* |
| --- | --- | --- |
|  | **N=80** |  |
| Age | 62.2 (14.0) | 80 |
| Sex: |  | 80 |
| Female | 35 (43.8%) |  |
| Male | 45 (56.2%) |  |
| Overall Survival Months | 26.7 (18.1) | 80 |
| Overall Survival Status: |  | 80 |
| Deceased | 23 (28.7%) |  |
| Living | 57 (71.2%) |  |
| Progress Free Survival Months | 23.2 (17.5) | 79 |
| Progression Free Status: |  | 80 |
| Censored | 50 (62.5%) |  |
| Progression | 30 (37.5%) |  |
| Disease specific Survival status: |  | 80 |
| Alive or Dead Tumor Free | 59 (73.8%) |  |
| Dead with Tumor | 21 (26.2%) |  |
| Recurrence: |  | 80 |
| Non-recurrent | 54 (67.5%) |  |
| Recurrent | 26 (32.5%) |  |
| Fraction Genome Altered | 0.16 (0.12) | 80 |
| Mutation Count | 16.9 (42.3) | 80 |
| SCNA cluster: |  | 80 |
| 1 | 15 (18.8%) |  |
| 2 | 23 (28.7%) |  |
| 3 | 22 (27.5%) |  |
| 4 | 20 (25.0%) |  |
| BAP1 mutation | 13 (16.2%) | 80 |
| Chromosome 3 status: |  | 80 |
| Disomy | 21 (26.2%) |  |
| Monosomy | 31 (38.8%) |  |
| 'Missing' | 28 (35.0%) |  |
| Chromosome 8 status: |  | 80 |
| Disomy | 19 (23.8%) |  |
| Polysomy | 33 (41.2%) |  |
| 'Missing' | 28 (35.0%) |  |
| Immune cluster: |  | 80 |
| High | 24 (30.0%) |  |
| Low | 56 (70.0%) |  |
| Mutation: |  | 80 |
| GNA11 p.Q209L | 34 (42.5%) |  |
| GNAQ p.Q209L | 9 (11.2%) |  |
| GNAQ p.Q209P | 27 (33.8%) |  |
| GNAQ,GNA11 p.Q209L,p.R166H | 1 (1.25%) |  |
| WT | 9 (11.2%) |  |
| prot: |  | 80 |
| p.G48V | 1 (1.25%) |  |
| p.Q209L | 44 (55.0%) |  |
| p.Q209P | 27 (33.8%) |  |
| p.R183Q | 1 (1.25%) |  |
| p.R183Q,p.R183C | 1 (1.25%) |  |
| WT | 6 (7.50%) |  |

**Supplementary Table 2.** Baseline characteristics of 71 mutated samples from TCGA-UM by amino-acid change. P-values for categorical variables were calculated by Chi-Squared Tests. P-values for continuous variables were calculated by Wilcoxon tests.

|  | P.Q209L<br>N=44 | P.Q209P<br>N=27 | P-VALUE |
| --- | --- | --- | --- |
| Age | 62.7 (14.7) | 62.1 (13.2) | 0.850 |
| Sex: |  |  | 0.399 |
| Female | 17 (38.6%) | 14 (51.9%) |  |
| Male | 27 (61.4%) | 13 (48.1%) |  |
| Overall Survival Months | 25.0 (16.7) | 27.9 (21.3) | 0.549 |
| Overall Survival Status: |  |  | 0.341 |
| Deceased | 14 (31.8%) | 5 (18.5%) |  |
| Living | 30 (68.2%) | 22 (81.5%) |  |
| Progress Free Survival Months | 21.1 (16.5) | 25.0 (19.9) | 0.405 |
| Progression Free Status: |  |  | 0.304 |
| Censored | 26 (59.1%) | 20 (74.1%) |  |
| Progression | 18 (40.9%) | 7 (25.9%) |  |
| Disease specific Survival status: |  |  | 0.260 |
| Alive or Dead Tumor Free | 31 (70.5%) | 23 (85.2%) |  |
| Dead with Tumor | 13 (29.5%) | 4 (14.8%) |  |
| Recurrence: |  |  | 0.062 |
| Non-recurrent | 27 (61.4%) | 23 (85.2%) |  |
| Recurrent | 17 (38.6%) | 4 (14.8%) |  |
| Fraction Genome Altered | 0.15 (0.10) | 0.14 (0.11) | 0.630 |
| Mutation Count | 11.1 (4.17) | 13.3 (3.77) | **0.028** |
| SCNA cluster: |  |  | 0.160 |
| 1 | 7 (15.9%) | 8 (29.6%) |  |
| 2 | 11 (25.0%) | 10 (37.0%) |  |
| 3 | 13 (29.5%) | 3 (11.1%) |  |
| 4 | 13 (29.5%) | 6 (22.2%) |  |
| BAP1 mutation | 7 (15.9%) | 5 (18.5%) | 0.757 |
| Chromosome 3 status: |  |  | 0.384 |
| Disomy | 11 (37.9%) | 9 (56.2%) |  |
| Monosomy | 18 (62.1%) | 7 (43.8%) |  |
| Chromosome 8 status: |  |  | 0.598 |
| Disomy | 9 (31.0%) | 7 (43.8%) |  |
| Polysomy | 20 (69.0%) | 9 (56.2%) |  |
| Immune cluster: |  |  | 1.000 |
| High | 13 (29.5%) | 8 (29.6%) |  |
| Low | 31 (70.5%) | 19 (70.4%) |  |

**Supplementary Table 3.**Peptides predicted to bind to HLA by MHCSeqNet, NetMHC, NetMHCpan, MHCflurry, MixMHCpred, NetMHCcons and NetMHCpanstab tools. Attached in excel format.

**Supplementary Table 4.**
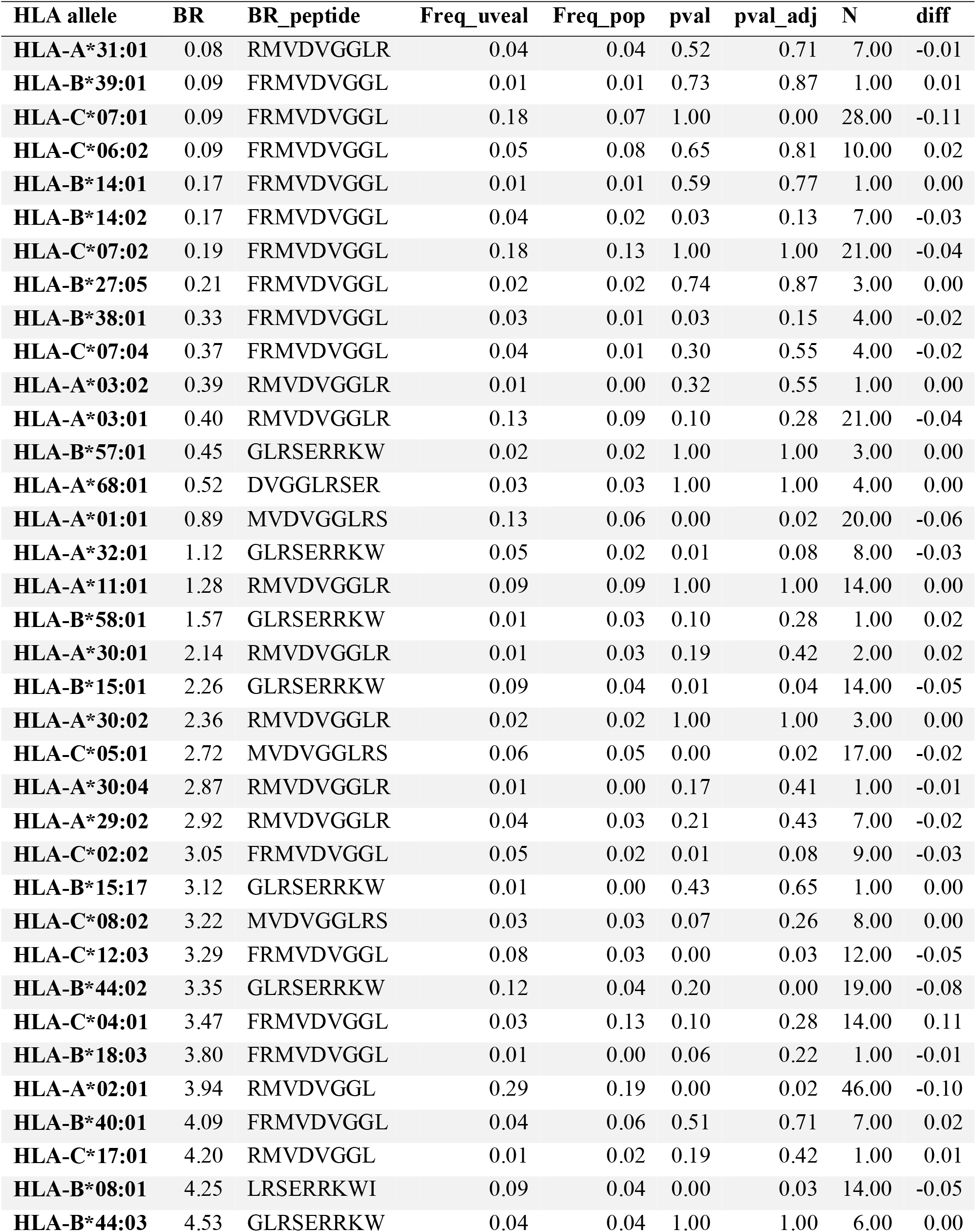
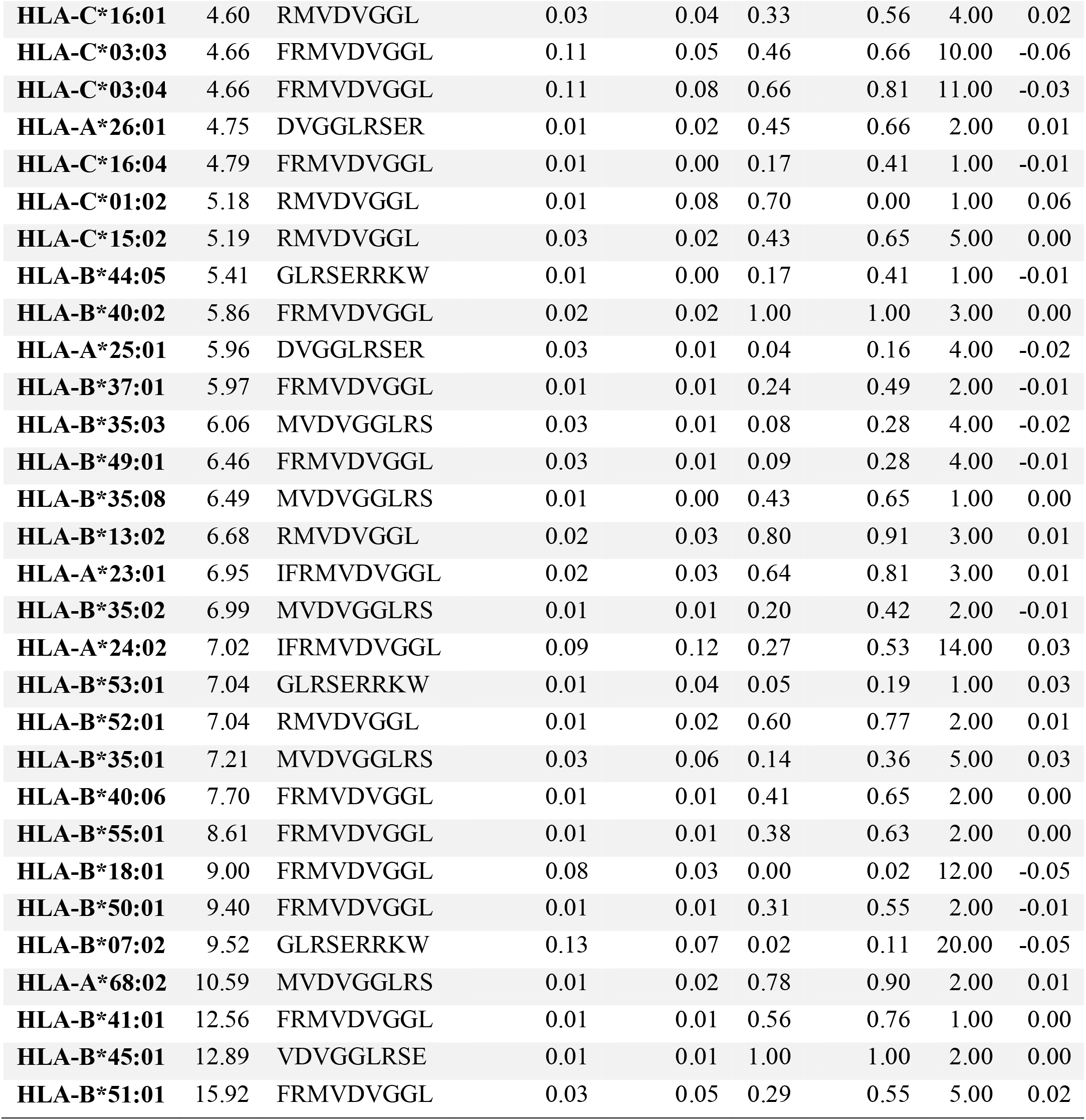
Results from bionomial tests comparing HLA frequencies between general population and uveal melanoma population, ordered by BR score. HLA haplotypes with statistical differences and higher frequency in UM are coloured in yellow. Only HLA haplotypes present in UM patients were compared. BR; Q209L BR score, BR_peptide; 9 mer amino acid of best rank for the allele. Freq_uveal; frequency of haplotype in uveal melanoma samples, Freq_pop; frequency of haplotype in the 1000Genomes population, pval_adj; p-value adjuted by FDR, N; number of UM patients with the HLA allele, diff; frequency difference.

**Supplementary Table 5.**Differentially expressed genes between GNAQ and GNA11 samples. Attached in excel.

**Supplementary Table 6.** Functional analysis results.

| Description | setSize | Enrichment Score | NES | pvalue | p.adjust | qvalue | rank |
| --- | --- | --- | --- | --- | --- | --- | --- |
| HALLMARK_INTERFERON_GAMMA_RESPONSE | 198 | 0.6380 | 1.6863 | 0.0000 | 0.0000 | 0.0000 | 4437 |
| HALLMARK_INTERFERON_ALPHA_RESPONSE | 96 | 0.6771 | 1.7315 | 0.0000 | 0.0000 | 0.0000 | 4437 |
| HALLMARK_MTORC1_SIGNALING | 197 | 0.5482 | 1.4486 | 0.0000 | 0.0001 | 0.0000 | 4769 |
| HALLMARK_ALLOGRAFT_REJECTION | 196 | 0.5373 | 1.4196 | 0.0000 | 0.0001 | 0.0001 | 4159 |
| HALLMARK_COMPLEMENT | 200 | 0.5136 | 1.3583 | 0.0002 | 0.0015 | 0.0009 | 4711 |
| HALLMARK_IL6_JAK_STAT3_SIGNALING | 87 | 0.5850 | 1.4823 | 0.0002 | 0.0015 | 0.0009 | 3108 |
| HALLMARK_INFLAMMATORY_RESPONSE | 200 | 0.5020 | 1.3278 | 0.0005 | 0.0036 | 0.0023 | 5102 |
| HALLMARK_PROTEIN_SECRETION | 96 | 0.5589 | 1.4292 | 0.0009 | 0.0053 | 0.0034 | 6435 |
| HALLMARK_COAGULATION | 138 | 0.5170 | 1.3438 | 0.0012 | 0.0069 | 0.0044 | 1783 |
| HALLMARK_TNFA_SIGNALING_VIA_NFKB | 199 | 0.4889 | 1.2927 | 0.0021 | 0.0103 | 0.0065 | 4152 |
| HALLMARK_APOPTOSIS | 159 | 0.5016 | 1.3130 | 0.0023 | 0.0105 | 0.0066 | 3642 |
| HALLMARK_HYPOXIA | 197 | 0.4856 | 1.2832 | 0.0029 | 0.0120 | 0.0076 | 3143 |
| HALLMARK_UNFOLDED_PROTEIN_RESPONSE | 110 | 0.5168 | 1.3288 | 0.0069 | 0.0266 | 0.0168 | 5050 |
| HALLMARK_OXIDATIVE_PHOSPHORYLATION | 200 | 0.4693 | 1.2412 | 0.0079 | 0.0281 | 0.0177 | 6606 |
| HALLMARK_FATTY_ACID_METABOLISM | 156 | 0.4796 | 1.2541 | 0.0149 | 0.0496 | 0.0313 | 6130 |

